# Single-cell molecular characterization to partition the human glioblastoma tumor microenvironment (TME) genetic background

**DOI:** 10.1101/2022.02.10.479701

**Authors:** Francesca Lessi, Sara Franceschi, Mariangela Morelli, Michele Menicagli, Francesco Pasqualetti, Orazio Santonocito, Carlo Gambacciani, Francesco Pieri, Filippo Aquila, Paolo Aretini, Chiara Maria Mazzanti

## Abstract

Glioblastoma (GB) is a devastating primary brain malignancy. Recurrence of GB is inevitable despite the standard treatment of surgery, chemotherapy, and radiation, and the median survival is limited to around 15 months. Barriers to treatment include the complex interactions among the different cellular components inhabitant of the tumor microenvironment. These challenges are further compounded by extensive inter- and intra-tumor heterogeneity and by the dynamic plasticity of GB cells. The complex heterogeneous nature of GB cells is helped by the local inflammatory tumor microenvironment, which mostly induces tumor aggressiveness and drug resistance. More effective therapy development heavily depends on higher resolution molecular subtype signatures. All cellular components of a GB tumor are subjected to a continuous pressure inducing proliferation that might favor the accumulation of genetic mutations. Understanding the genetic background of several cellular components belonging to a GB microenvironment could give insights into tumor behavior and progression. In the present study by using fluorescent multiple labeling and DEPArray cell separator, we recovered several single cells or groups of single cells from populations of different origins from IDH-WT GB samples. From each GB sample, we collected astrocytes (GFAP+), microglia (IBA+), stem cells (CD133+), and endothelial cells (CD105+) and performed Copy Number Aberration (CNA) analysis with a low sequencing depth. The same tumors were subjected to a bulk CNA analysis. The tumor partition in its single components allowed single-cell molecular subtyping which revealed new aspects of GB altered genetic background. Nowadays, single-cell approaches are leading to a new understanding of GB physiology and disease. Moreover, single-cell CNAs resource will permit new insights into genome heterogeneity, mutational processes, and clonal evolution in malignant tissues.

## 1 Introduction

Glioblastoma (GB) is the most aggressive and deadly primary tumor of the central nervous system in adults with overall survival of fewer than 15 months (1). The GB impressive poor prognosis, despite the development in recent decades of new and innovative therapies, is enhanced by the resistance developed towards radio and chemotherapy (2). In this tumor, as well as in other cancer types, the tumor microenvironment (TME) plays a pivotal role in treatment resistance (2). GB microenvironment is composed of a massive of different cells, and besides malignant astrocytes and cancer stem cells, stromal, endothelial cells, pericytes and a huge number of immune cells are present (3). Moreover, intratumoral heterogeneity (ITH), which is one of the major features of GB tumor, is also hugely involved in anticancer treatments resistance (4)(5) and is critical to promote tumoral growth and aggressiveness (6). In support of this last remark, it has recently been demonstrated in GB that within the same tumor co-exist different sub-clones that respond differently to the drug therapy (7). These sub-populations of cells show distinct genomic profiles that reveal an individual behavior peculiar from the whole cell population (8). Indeed currently, the single-cell approach in GB is becoming increasingly popular. The single-cell isolation allows the selection of a specific cell population that comprises <1% of the total cells (9,10). Reaching single-cell resolution enables avoiding the averaging of bulk analysis and capturing the heterogeneity of cells. Copy number aberration (CNA) is one of the most important somatic alterations in cancer (11) (12) meant as somatic changes to chromosome structure such as gain and deletion of a particular DNA segment (> 1 kb) (13). The most common CNAs in GB include loss, or partial loss, of chromosomes 9 and 10; gain of chromosomes 7, 19, and 20; focal deletion of CDKN2A/B locus (9p21.3); and focal high-level amplification of EGFR locus (7p11.2) (14) (16). In particular, it is well known that CNAs targeting chromosomes 7 and 10 are some of the earliest events in GB tumor evolution. The analysis of these aberrations is interesting because CNAs are detected with much greater accuracy than individual mutations and are associated with ITH in most cancers. Moreover, the aggregation of cells sharing the same CNA profiles allows improving the phylogenetic analysis at single nucleotide levels (16).

In this work, we collected three human GB tumors and after dissociation, a single-cell molecular characterization was carried out, with particular attention on four cell populations: astrocytes, microglia cells, endothelial cells, and stem cells. We collected a certain number of single and groups of single cells belonging to the populations mentioned above using a high throughput selective sorting technology and we investigated the genomic aberrations (CNA analysis) in these different types of tumor cells. The whole parental tumors were subjected to a bulk CNA analysis, as well, to compare their molecular profile with single cell results. The tumor partition in its single components allowed single-cell molecular profiling which revealed new aspects of GB altered genetic background. Our work demonstrates that the single-cell approach is more representative and detailed than the bulk analysis, which contributes to a deeper insight into the basic molecular mechanisms of glioblastoma.

## 2 Materials and Methods

### 2.1 Human glioblastoma tissue collection

The study has been performed according to the declaration of Helsinki and the sample’s collection protocol was approved by the Ethics Committee of the University Hospital of Pisa (787/2015). Tumor tissues were obtained from patients who underwent surgical resection of histologically confirmed GB after informed consent. Samples were obtained from the Unit of Neurosurgery of Livorno Civil Hospital. Three patient cases (GB01, GB02 and GB03) were included in the present study, the clinical and demographic data and the pathological and therapeutical information are summarized in Table 1. All cases had a diagnosis of GB with no previous history of any brain neoplasia and were not carrying R132 IDH1 or R172 IDH2 mutations.

**Table 1.**
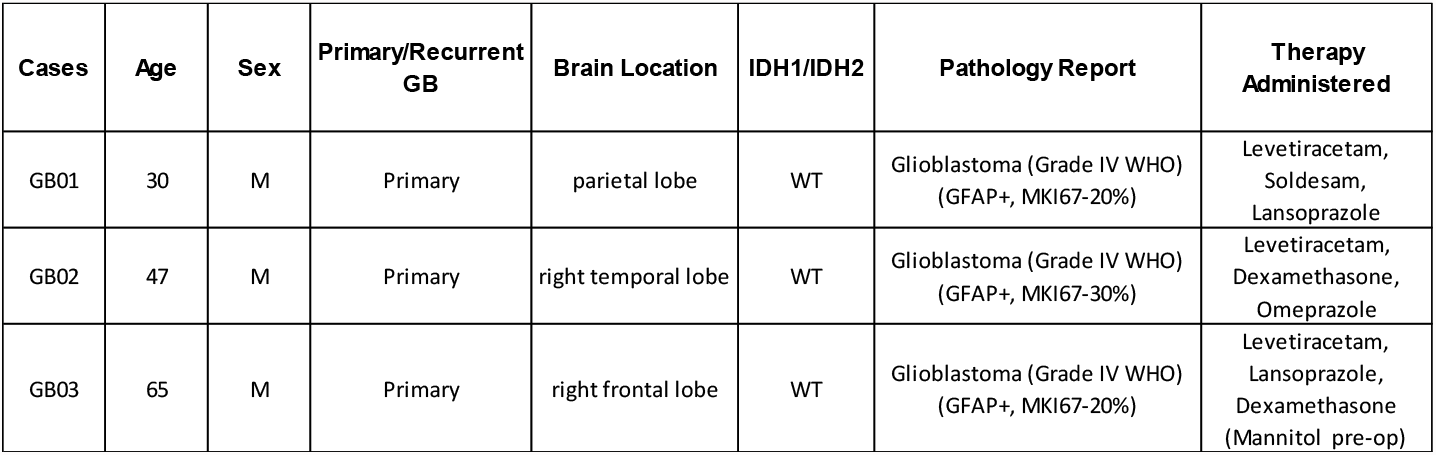
Patient clinical, demographic, pathological, and therapeutical data.

Surgically resected tumors were collected and stored in MACS tissue storage solution (Miltenyi Biotec, Bergisch Gladbach, Germany) at 4°C for 2–4 hours. Each tumor sample was washed with Dulbecco’s phosphate-buffered saline (DPBS) in a sterile dish and portioned with a scalpel into about 0.5-2 cm^2^ pieces under a biological hood. Afterward, they were vital frozen at -140°C in 90% fetal bovine serum (FBS) and 1% dimethyl sulfoxide (DMSO) for further analyses.

### 2.2 Tumor dissociation to single-cell suspensions

Frozen GB tissues were defrosted in a water bath at 37°C, were washed with DPBS in a sterile dish and cut with a scalpel into small pieces. These finely minced tumor chunks were transferred in a C-tube (Miltenyi Biotec) with the appropriate volume of buffer X following the protocol (Brain Tumor Dissociation Kit, Miltenyi Biotec) for the tumor dissociation with the gentleMACs Dissociator (Miltenyi Biotec).

### 2.3 Immunofluorescence of single-cell suspensions

The cell suspensions obtained were transferred in 1.5 ml LoBind tubes and washed three times with DPBS. After centrifugation at 300g for 10 minutes at room temperature, the supernatant was removed and the cells resuspended with 400 µl of Running Buffer composed of MACS BSA stock solution (Miltenyi Biotec) 1:20 with autoMACS Rinsing Solution (Miltenyi Biotec). The cells were fixed adding 400 µl of Paraformaldehyde 4%. Fixation solution was incubated for 20 minutes at room temperature. To stop the reaction, the sample tubes were filled with DPBS and centrifuged at 400g for 5 minutes at room temperature. Afterward, we performed two washes with DPBS to the sample tubes and then we incubated the pellet with blocking solution for 10 minutes at room temperature (BSA 3% in DPBS). The blocking reaction was stopped filling the tube with DPBS and centrifuged at 400g for 5 minutes at room temperature. The cells were resuspended in Running Buffer and counted with Luna Automated Cell Counter (Logos Biosystems, Gyeonggi-do 14055, South Korea). For the immunofluorescence, a maximum of 100.000 fixed cells was used for the staining. The antibodies chosen for the staining were: anti-GFAP APC (130-124-040, Miltenyi Biotec) for astrocytes, anti-Iba1 PE (ab209942, Abcam, Cambridge, UK) for microglia cells, anti-CD105 PerCP/Cy5.5 (ab234265, Abcam, Cambridge, UK) for endothelial cells, anti-CD133 FITC (11-1339-42, eBioscience, San Diego, CA, USA) for stem cells and Hoechst 33342 (62249, Thermofisher Scientific, Waltham, MA, USA) for nuclei. Twenty µl of anti-CD105 and 25 µl of anti-CD133 were added to the cell suspensions and mixed by gently pipetting. The samples were incubated for 15 minutes in the dark at 4 °C. The reaction was stopped by adding 1 ml of Running Buffer and mixed by gently pipetting. Then the sample tubes were centrifuged at 400g for 10 minutes at room temperature, the supernatant was removed and the cells were resuspended with 100 µl of Inside Perm Buffer (Inside Stain Kit, Miltenyi Biotec). Eight µl of anti-GFAP and 2.5 µl of anti-Iba1 were added to the cell suspensions and mixed by gently pipetting. The samples were incubated for 20 minutes in the dark at room temperature. The reaction was stopped by adding 1 ml of Inside Perm Buffer and mixed by gently pipetting. Then the sample tubes were centrifuged at 400g for 10 minutes at room temperature, the supernatant was removed and the cells were resuspended with 1 ml of Running Buffer. One µl of Hoechst (1mg/ml) was added to the sample tubes and mixed by gently pipetting. The samples were incubated for 5 minutes in the dark at room temperature. Then the sample tubes were centrifuged at 400g for 10 minutes at room temperature and resuspended in 200 µl of Running Buffer.

### 2.4 Single cell Isolation by DEPArray™ NxT

Single cells were isolated and sorted with DEPArray NxT (Menarini, Silicon Biosystems, Bologna, Italy). After the immunofluorescence of the single cell suspensions, the cells were counted, we used a maximum of 24.000 cells to load the DEPArray NxT Cartridge. The samples were washed two times with 1 ml of SB115 Buffer (Menarini, Silicon Biosystems) and the cells were loaded on the DEPArray NxT cartridge following the protocol instructions. CellBrowser™ analysis software, integrated into the DEPArray™ system, allows to view and to select cells from the particle database according to multiple criteria, based on qualitative and quantitative marker evaluation and cell morphology. This software enables to create populations and sub-populations of cells using some analysis tools as scatter plots, histograms, and image panels. Cells become un-routable based on their positions, when these are out of the cage it is no longer possible to move them and therefore complete the recovery. First of all, we excluded clusters of two or three cells, clumps, and spurious events and focused only on single cells with the desired fluorescence analyzing only the “centered” DAPI cells in the cage. The single cells were selected manually based on fluorescence labeling and morphology. About 20 different single cells were recovered for each tumor patient and volume reduction was performed with VRNxT-Volume Reduction Instrument (Menarini, Silicon Biosystems) according to the instruction manual. The isolated cells were stored at −20 °C until later downstream analyses.

### 2.5 DNA Extraction from fresh tissues

Genomic DNA was extracted directly from up to 50 mg of fresh tissue of GB01, GB02 and GB03 using the Maxwell® 16 Instrument with the Maxwell® 16 Tissue DNA Purification Kit (Promega, Madison, WI). DNA concentration was determined using the Qubit Fluorometer (Life Technologies, Carlsbad, CA) and the quality was assessed using the Agilent 2200 Tapestation (Agilent Technologies, Santa Clara, CA) system.

### 2.6 Ampli1™ whole genome amplification and Low Pass Analysis

Whole-genome amplification on all recovered single cells was performed using the Ampli1**™** WGA Kit version 02 (Menarini, Silicon Biosystems) following the manufacturer’s instructions. The same procedure was adjusted for the DNA obtained from fresh tissues starting from 1 µl of 1 ng/ µl. Afterward, the WGA product was cleaned up with SPRIselect Beads (Beckman Coulter, Brea, CA, USA) and sequencing-ready libraries were prepared with Ampli **™** LowPass Kit (Menarini, Silicon Biosystems) to detect chromosomal aneuploidies and copy number aberrations (CNAs) with a low sequencing depth. To sequence our libraries, we used Ion 520/530-OT2 kit (Ion Torrent, Life Technologies, Grand Island, NY) with the Ion 530 Chip (Ion Torrent). The runs were run on the Ion S5 system (Ion Torrent).

### 2.7 CNA calling

The data obtained from low-pass whole genome sequencing were processed with IchorCNA tool (17). The CNV segmented number profiles obtained from IchorCNA were processed with the CNApp tool (18) with default cutoffs.

## 3 Results

### 3.1 Isolation of single-cells from GB fresh tissues with DEPArray ™ NxT

Three GB fresh tissues obtained from the Unit of Neurosurgery of Livorno Civil Hospital were analyzed with DEPArray™ NxT, the overview of the procedure is shown in Figure 1 in which also H&E images for each tumor tissue are present. After the DEPArray NxT Cartridge loading, we selected the routable cells using the CellBrowser™ analysis software. In detail, for GB01, 2880 routable cells, for GB02, 17378 routable cells, and for GB03, 4788 routable cells, were observed. After that, we performed the exclusion of cell clusters obtaining single and routable cells: 2654, 9535 and 4278 cells respectively for GB01, GB02 and GB03. Afterward, we identified four main populations (astrocytes, microglia cells, endothelial cells and stem cells) and several cells with double fluorescence staining. An example of the main populations is shown in Figure 2. In Figure 3, percentages of the main populations, found in the 3 samples, are summarized, while in Supplementary Materials (Figure 1S) double fluorescence stained cells and no labeled cells are shown. We recovered both single cells and groups of a maximum of 5 single cells with the same characteristics.

**Figure 1.**
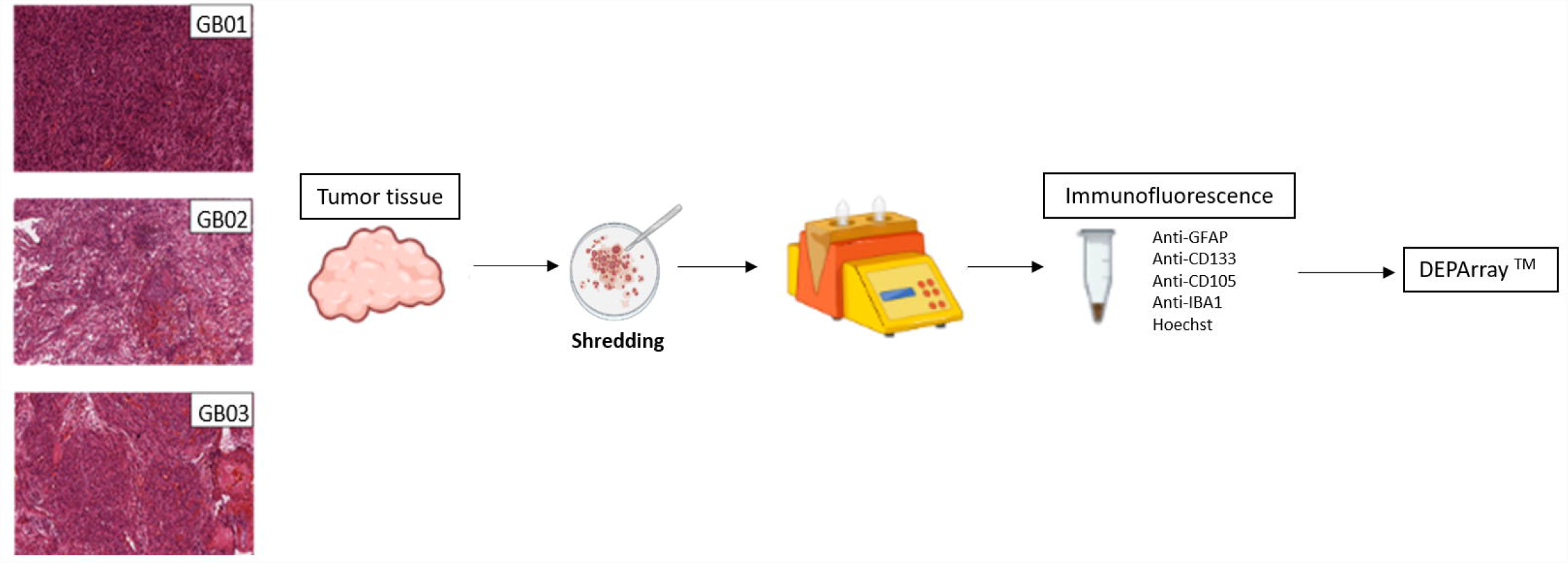
Histological images of GB01, GB02 and GB03. Experimental design starting from tumor shredding to DEPArray analysis.

**Figure 2.**
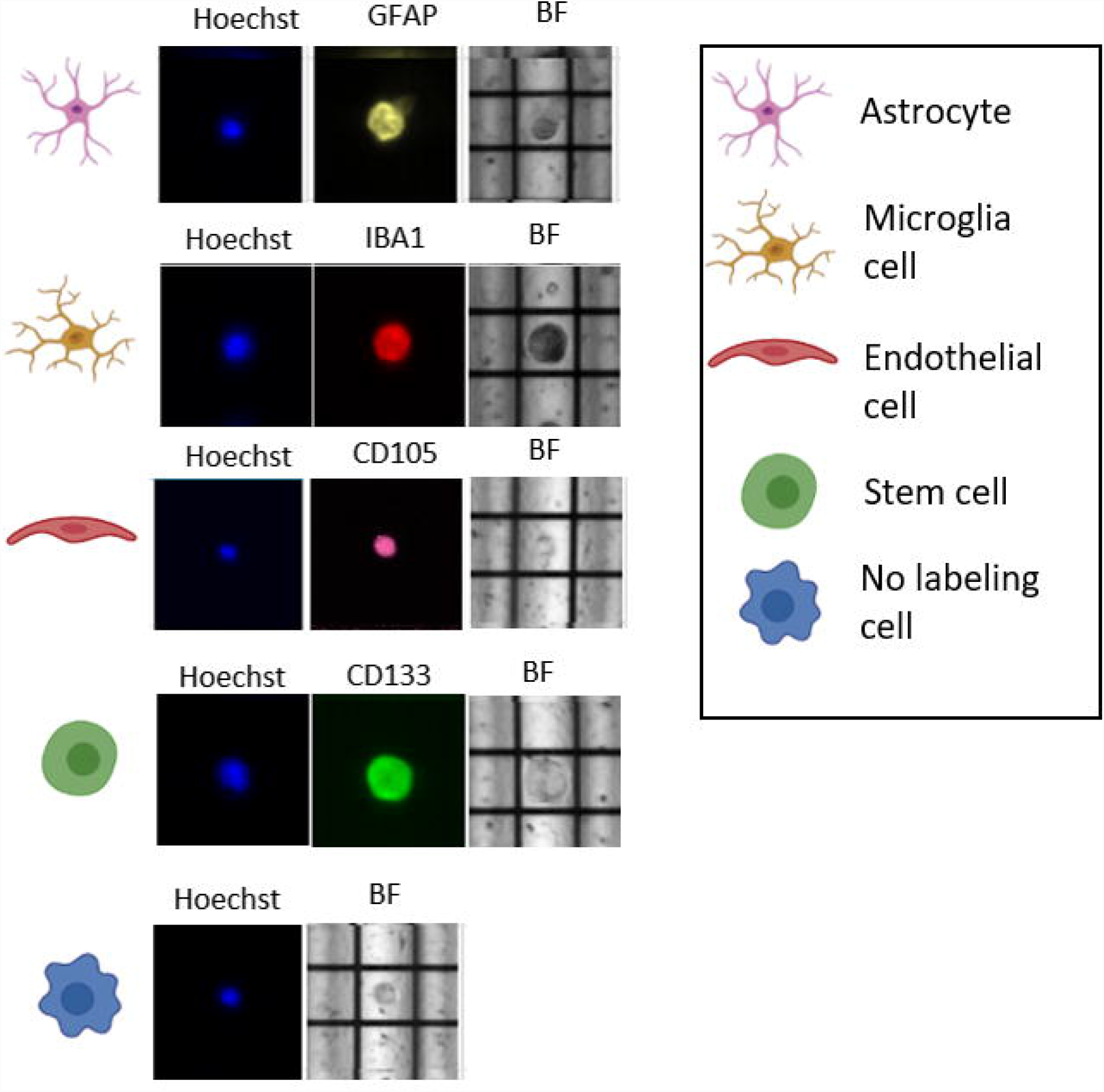
Example of DEPArray images of single cells belonging to the main GB populations, stained in yellow with GFAP (astrocytes), in red with IBA1 (microglia), in purple with CD105 (endothelial cells), in green with CD133 (stem cells) and in blue with Hoechst. BF: Brightfield.

**Figure 3.**
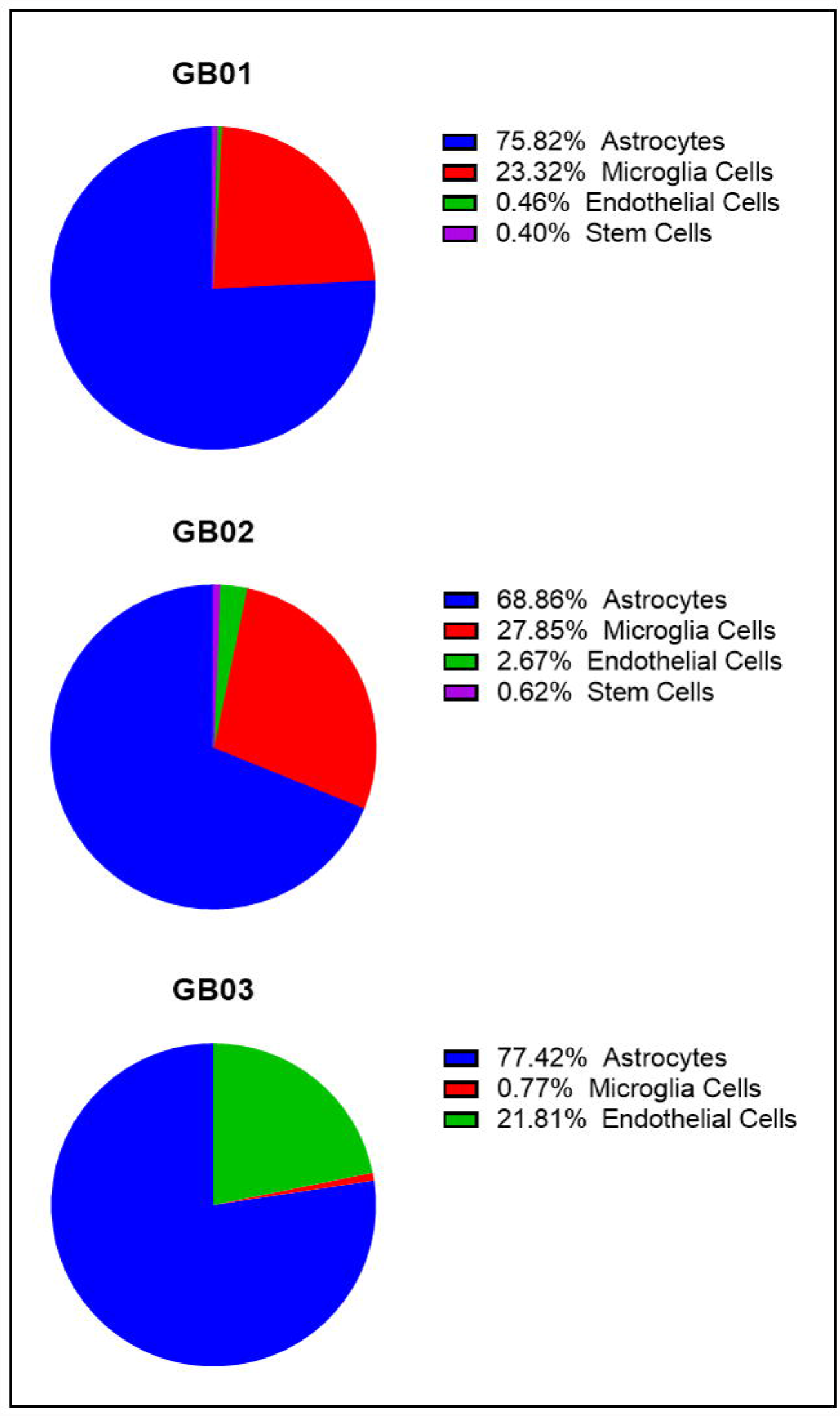
Pie charts of the percentages of the main cell type populations found in GB01, GB02 and GB03.

Picked cells for the 3 samples are summarized in Figure 4. In particular, for GB01 we selected 20 cells: 3 single astrocytes, 3 groups of astrocytes, 4 microglia single cells, 2 groups of microglia cells, 1 group of endothelial cells, 1 single stem cell, 2 single astrocytes/microglia cells (positive both for GFAP and IBA1), 3 single cells and 1 group of single cells without labeling (positive to Hoechst 33342 only). For GB02, recovered cells were 26: 6 single astrocytes, 5 microglia single cells, 5 single endothelial cells, 3 groups of endothelial cells, 5 single stem/endothelial cells (positive both for CD133 and CD105), 1 group of stem/endothelial cells (positive both for CD133 and CD105) and 1 single cell without labeling (only Hoechst 33342 signal). Finally, for GB03 the cells selected were 17: 6 single astrocytes, 5 single microglia cells and 6 single endothelial cells.

**Figure 4.**
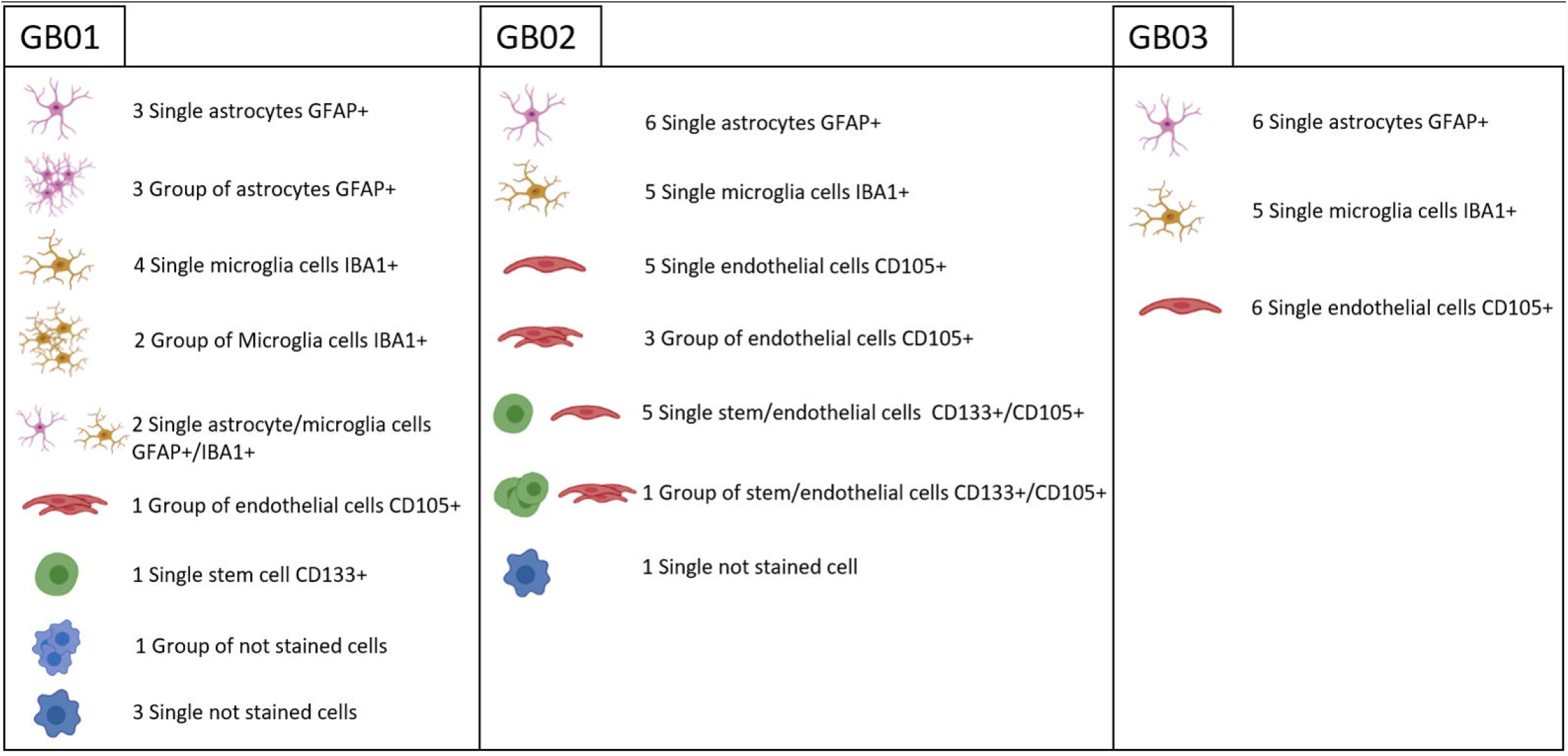
Summary of all the recovered cells after DEPArray analysis from GB01, GB02 and GB03. Astrocytes, microglia, endothelial and stem cells were collected. Double staining cells and only Hoechst positive cells are shown.

### 3.2 Copy Number Aberrations (CNAs) Analysis

The cellular genomic profiling was performed on selected cells using the Ampli1™ LowPass kit to identify genome-wide CNAs at the single-cell level and to obtain information on ITH. The same analysis was carried out also on DNA obtained from tumor fresh tissues (GB01, GB02 and GB03), to compare the bulk molecular profile to the one derived from single cells.

In Figure 5 the CNA pattern of the fresh GB tissues is shown: as expected, each sample has a different CNA configuration due to GB ITH. However, all three samples presented chromosome 10q deletion and GB01 and GB02 also chromosome 7 amplification, which represents typical GB alterations. Consequently, for each sample, tumors in bulk and single cells CNAs were compared. The description of these results is summarized in Tables 2-3-4.

**Table 2.** CNAs results obtained after CNApp processing for single cells and groups of single cells collected in the GB01 sample.

| **GB01** |  |
| --- | --- |
| **Single cells collected** | **CNA** |
| Group of endothelial cells | WT |
| Microglia cell | 19- |
| Microglia cell | WT |
| Microglia cell | 1p-, 7+, 9q+, 10-, 17q+, 19+ |
| Microglia cell | 1q+, 2-, 5+, 7+, 9p-, 9q+, 10-, 17q+, 19+ |
| Group of microglia cells | WT |
| Group of microglia cells | 7+, 9q+, 10-, 17q+ |
| Astrocyte | 1p-, 7+, 9q+, 10-, 17q+, 19+ |
| Astrocyte | 1p-, 7+, 9q+, 10-, 17q+, 19+ |
| Astrocyte | 1q+, 5+, 7+, 9p-, 9q+, 10-, 17p-, 17q+, 19q+ |
| Group of astrocytes | 1p-, 7+, 9q+, 10-, 17p-, 17q+, 19+ |
| Group of astrocytes | 1p-, 7+, 9+, 10-, 17p-, 17q+, 19+ |
| Group of astrocytes | 1p-, 7+, 9q+, 10-, 17q+ |
| Stem cell | 1p-, 7+, 9+, 10-, 17q+, 19q+ |
| Astrocyte/microglia cell | 1p-, 7+, 9+, 10-, 17q+, 19q+ |
| Astrocyte/microglia cell | 1p-, 7+, 9+, 10-, 17q+, 19+ |
| Not stained cell | WT |
| Not stained cell | WT |
| Not stained cell | 1p-, 7+, 9q+, 10-, 17p-, 17q+, 19q+ |
| Group of not stained cells | WT |

**Table 3.**
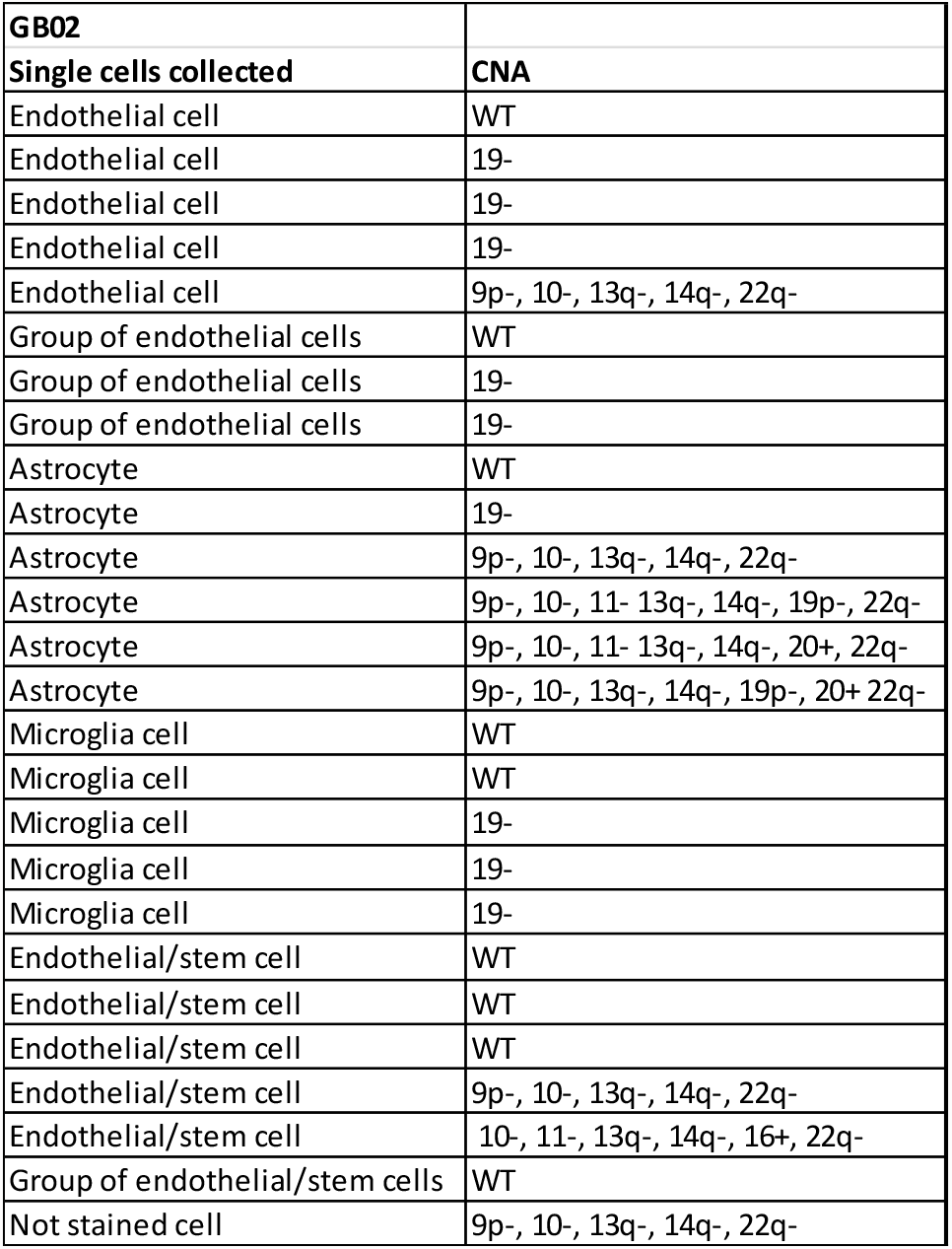
CNAs results obtained after CNApp processing for single cells and groups of single cells collected in the GB02 sample.

| **GB02** |  |
| --- | --- |
| **Single cells collected** | **CNA** |
| Endothelial cell | WT |
| Endothelial cell | 19- |
| Endothelial cell | 19- |
| Endothelial cell | 19- |
| Endothelial cell | 9p-, 10-, 13q-, 14q-, 22q- |
| Group of endothelial cells | WT |
| Group of endothelial cells | 19- |
| Group of endothelial cells | 19- |
| Astrocyte | WT |
| Astrocyte | 19- |
| Astrocyte | 9p-, 10-, 13q-, 14q-, 22q- |
| Astrocyte | 9p-, 10-, 11-, 13q-, 14q-, 19p-, 22q- |
| Astrocyte | 9p-, 10-, 11-, 13q-, 14q-, 20+, 22q- |
| Astrocyte | 9p-, 10-, 13q-, 14q-, 19p-, 20+ 22q- |
| Microglia cell | WT |
| Microglia cell | WT |
| Microglia cell | 19- |
| Microglia cell | 19- |
| Microglia cell | 19- |
| Endothelial/stem cell | WT |
| Endothelial/stem cell | WT |
| Endothelial/stem cell | WT |
| Endothelial/stem cell | 9p-, 10-, 13q-, 14q-, 22q- |
| Endothelial/stem cell | 10-, 11-, 13q-, 14q-, 16+, 22q- |
| Group of endothelial/stem cells | WT |
| Not stained cell | 9p-, 10-, 13q-, 14q-, 22q- |

**Table 4.** CNAs results obtained after CNApp processing for single cells and groups of single cells collected in the GB03 sample.

|  |  |
| --- | --- |
| **GB03** |  |
| **Single cells collected** | **CNA** |
| Endothelial cell | WT |
| Endothelial cell | WT |
| Endothelial cell | WT |
| Endothelial cell | WT |
| Endothelial cell | 7+, 9p-, 9q+, 10-, 20+, 22q- |
| Endothelial cell | 3q-, 7+, 9p-, 9q+, 10-, 20+, 22q- |
| Microglia cell | WT |
| Microglia cell | WT |
| Microglia cell | WT |
| Microglia cell | WT |
| Microglia cell | WT |
| Astrocyte | WT |
| Astrocyte | 7+, 9p-, 9q+, 10-, 20+, 22q- |
| Astrocyte | 7+, 9p-, 9q+, 10-, 20+, 22q- |
| Astrocyte | 7+, 9p-, 9q+, 10-, 20+, 22q- |
| Astrocyte | 7+, 9p-, 9q+, 10-, 20+, 22q- |
| Astrocyte | 7+, 9p-, 9q+, 10-, 20+, 22q- |

**Figure 5.**
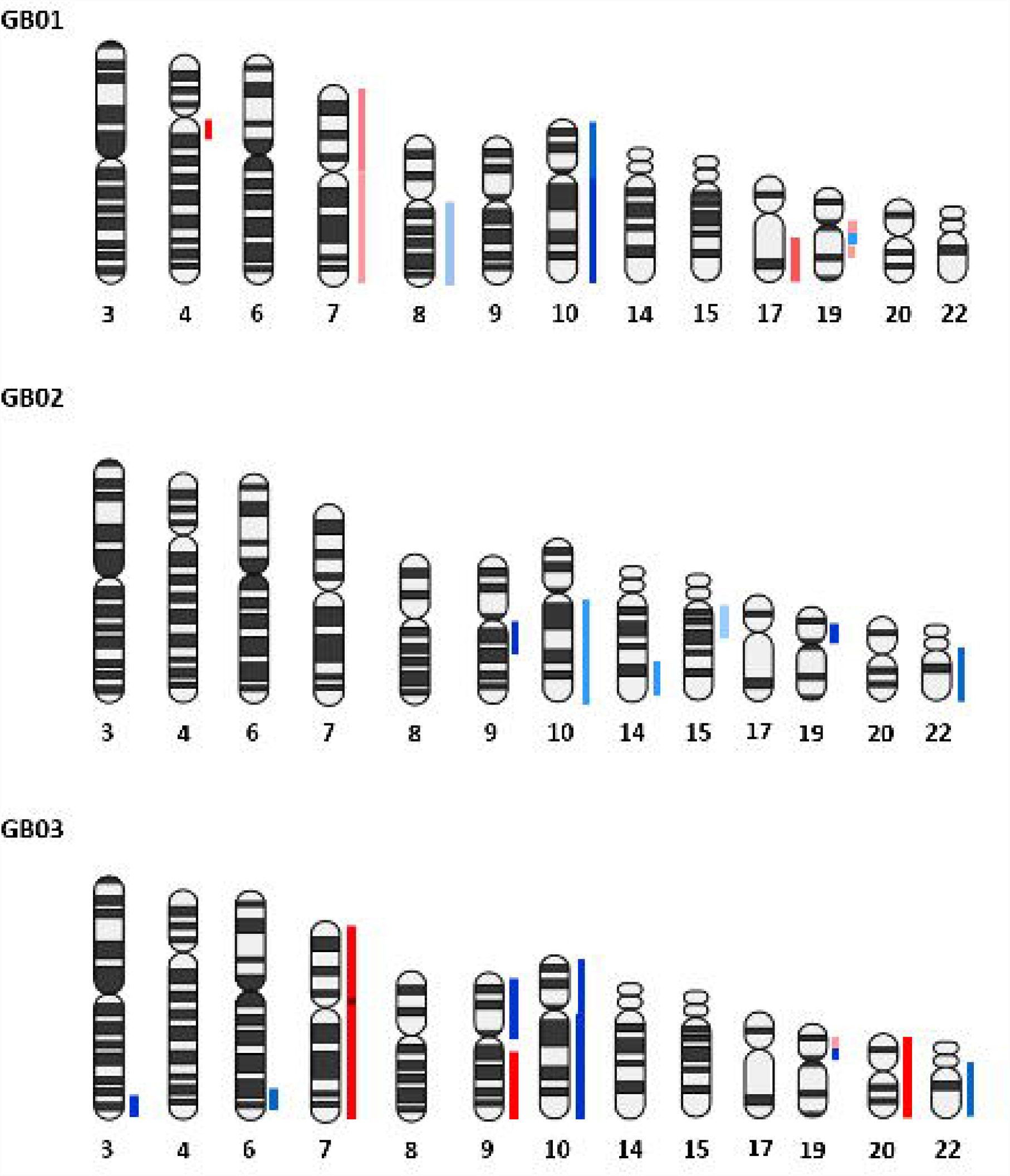
CNA pattern of the fresh GB tissues in bulk. The chromosomal amplifications are shown in red and in blue the deletions. The intensity of red and blue color components correlates to the gain and loss values based on the results obtained from the CNApp tool.

In GB01, we found a group of wild type endothelial cells; of 6 microglia cells (4 single cells and 2 groups of cells) 2 were wild type (one single cell and one group of cells), 1 cell showed chr 19 deletion only and the other cells showed different alterations sharing chr 10 deletion, and chr 7, 9q and 17q amplification; 6 astrocytes (3 single cells and 3 groups of cells) were altered sharing chr 10 deletion, and chr 7, 9q and 17q amplification; one stem cell with chr 1p and 10 deletion and chr 7, 9, 17q and 19q amplification. In GB01, moreover, 2 cells with double staining (GFAP and IBA1) were found with the same alterations, chr 1p and 10 deletions and chr 7, 9, 17q and 19q amplification. Finally, 3 out of 4 not stained cells (3 single and 1 group of cells) were wild type and 1 with chr 1p, 10 and 17p deletion and chr 7, 9q, 17q and 19q amplification.

In GB02 we found 8 endothelial cells (5 single and 3 groups of cells), 2 were wild type and the others carried chr 19 deletion except for only one having chr 9p, 10, 13q, 14q and 22q deletion. Then of 6 single astrocytes, 1 was wild type, 1 had chr 19 deletion, and the others shared chr 9p, 10, 13q, 14q and 22q deletion. Indeed, of 5 single microglia cells, 2 were wild type and 3 had chr 19 deletion. In GB02 we selected 6 double staining cells (CD133 and CD105 positive), of these, 4 were wild type and the others shared chr 10, 13q, 14q and 22q deletion. Finally, one not stained cell was wild type. GB03 counted 6 single endothelial cells, 4 of which were wild type and the other 2 presented different alterations sharing in particular chr 9p, 10 and 22q deletion and chr 7, 9q and 20 amplification. Five single microglia cells were all wild type. Finally, 6 single astrocytes were selected, 1 was wild type while the other cells showed all the same alterations: chr 9p, 10 e 22q deletion and chr 7, 9q and 20 amplification.

In Figure 6 the comparison between bulk fresh tumor CNAs and single-cells CNAs obtained with CNApp are shown.

**Figure 6.**
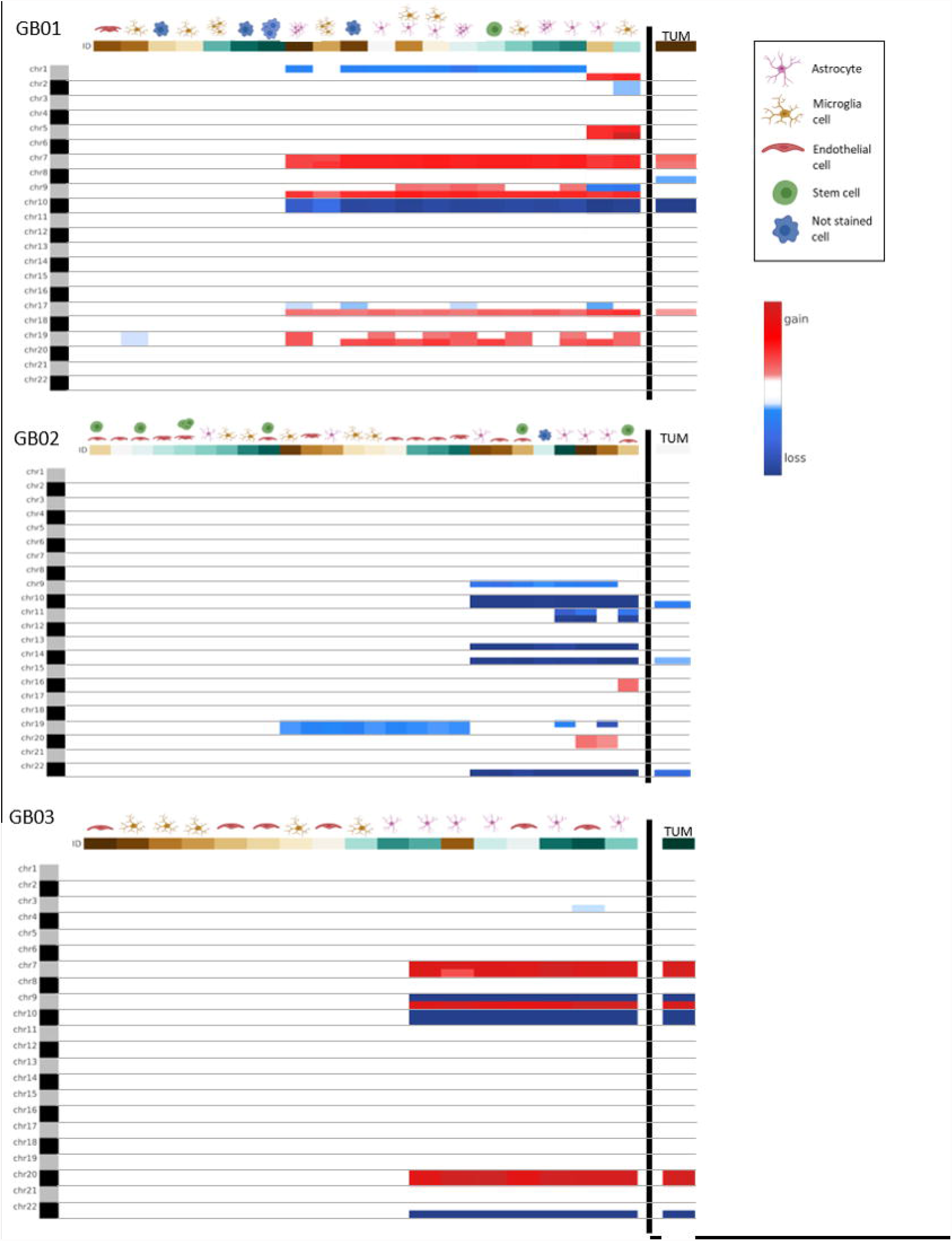
Genome-wide chromosome arm CNA profile heatmap for GB01, GB02 and GB03. For each sample the CNA profile of the single cells collected is shown and on the right the CNA profile of the tumor tissue in bulk.

## Discussion

Despite the new therapies developed in the last few years, GB still remains an incurable and devastating disease (19). The adjective *“multiforme*”, often used to define GB, was coined in 1926 by Percival Bailey and Harvey Cushing (20) to describe the various appearances of necrosis, cysts and hemorrhage. As a matter of fact, this definition also fits from a molecular point of view to explain the high degree of heterogeneity in GB. The poor prognosis of GB patients is mainly associated with ITH, which represents the presence in the tumor mass of multiple sub-clones, each characterized by different mutations (21). The sub-clones mutations are certainly masked during bulk tumor analysis (22). Nowadays, the single-cell analysis approach allows reading the DNA one cell at a time identifying the individual mutations that occurred in tumor progression. Recently, in some single-cell sequencing studies, to investigate the ITH, CNAs investigations were conducted instead of the identification of individual mutations with a gain in sensitivity and accuracy (4,23)(24).

In this work, we decided to focus our attention on some of the most iconic GB populations such as astrocytes, microglia, stem and endothelial cells to observe their molecular alterations and to compare them to the whole tumor tissue, in terms of CNAs.

Astrocytes are the star-shaped cells of the brain with different active roles both in the healthy and in brain pathological conditions (25). For example, they regulate neural signaling and give support in the blood-brain barrier (BBB) formation (25). Regarding GB tumorigenesis, a much-debated topic concerns the cell-of-origin in the cancer stem cell (CSC) or hierarchical hypothesis: GB stem cells (GSCs) or glioma initiating cells seem to be responsible for tumor formation (3). They are small population of stem cells characterized by self-renewal and differentiation properties (26). GSCs are involved in tumor growth, invasion and recurrence development (27). Based on this theory, GSCs can arise from neural stem cells (28) but also from already differentiated astrocytes transformed through genetic and epigenetic mutations (29)(30). Therefore, based on this hypothesis, initiating GB is composed of a mixture of cells including astrocytes and stem cells. In our work, the astrocytes, in general, in all the three tumors, were altered with a CNA pattern identical to the bulk tumor but in some cells with even more alterations, in support of the concept of the more sensitivity and accuracy of the single-cell analysis approach. Indeed, the only stem cell collected in GB02 showed a CNA pattern typical of a transformed tumor cell. This suggests that the cumulative acquisition of mutations in the stem cells can be responsible for invasive cancer generation.

In the brain, microglia cells represent the resident innate immune cells (macrophages) and these are involved in many crucial physiological processes (31). Microglia have been ignored for a long time but by now it is common knowledge that these cells are an integral part of the tumor, constituting approximately 30% of tumor mass (32) and participating in tumor progression and anti-cancer treatment resistance (33). Microglia cells have, indeed, a key role in many brain diseases (34). From our results, we observed some microglia cells with normal chromosomes sets, as we expected, but we also found some cells presenting CNAs, indicating that within the tumor there are also microglia cells with a potential tumoral behavior. From a transcriptional point of view some alterations have been described in GB microglia (35). In 2020, Maas and colleagues defined a particular type of transformed microglia cells. In this context tumoral GB cells hijack the microglia gene expression to enhance tumor proliferation suppressing the immune response (36).

Endothelial cells (ECs) represent the principal components of the BBB (37). Different brain pathologies, including GB, show molecular alterations of ECs (38). In GB, vessels are necessary for cancer cell spreading and it has been demonstrated that ECs regulate tumor invasion through crosstalk with GB cells (39). Our results illustrate the presence of wild type ECs but carrying also CNAs, confirming that in the tumor mass can be present tumor-ECs (also defined tumor-associated ECs) as it has been highlighted in some recent publications (40) (41,42). In these papers, the tumor-associated ECs showed different phenotypic and functional characteristics concerning normal ECs. Moreover, the relationship between ECs and GB tumor cells was demonstrated in two recent studies, in particular it was observed that tumor-derived ECs and GB stem cells shared the same genomic mutations and that CD144 and VDGFR2 genes are expressed by the emerging endothelium (43,44). Moreover, in our study, we observed and then recovered some cells with a double signal of labeling: astrocytes/microglia cells in GB01 and stem/endothelial cells in GB02. Indeed, in the literature has been reported the detection of dual positive cells in experiments using our same technology, especially in the circulating tumor cells studies (45). Fais *et al*. in 2007 introduced the concept of cannibalism as an exclusive property of malignant tumor cells (46). Moreover, Coopman *et al*. assumed that phagocytosis is the mechanism used by invasive tumor cells to allow migration in the surrounding tissues (47). In this regard, in malignant gliomas, phagocytic tumor cells were detected and particularly in GB (48) (49). A different hypothesis could be the cell fusion formation, for example, Huysentruyt *et al*. observed fusion between macrophages and tumor cells (50).

A further aspect that emerged from our results is the detection both in GB01 and in GB02 of some not-stained cells with CNAs. We observed, in fact, that not all the astrocytes are positive for GFAP and it has also been demonstrated in the literature that GFAP is not an astrocytes exclusive marker, as GFAP expression in GB varies significantly (51).

The use of CNAs as a method of evaluating tumor cells is more popular lately. CNA burden is assessed in different tumors, such as in prostate cancer (52) meaning as the analysis on variable amounts of amplifications or deletions in different patients. In particular, Hieronymus *et al*. (52) observed that patients with a high CNA burden showed a greater risk of relapse after treatment. For this reason, CNA analysis can be considered also as a useful marker. Therefore, tumor CNA burden, rather than individual CNAs, can be associated with cancer outcomes. Recently, CNAs analysis has been evaluated more advantageously than mutational analysis for diagnostic reasons in particular in association with survival (53): CNAs and miRNA analysis had a better performance rather than mutational data for poorly predicted survival. In addition, in melanoma, Roh *et al*. demonstrated that the association of CNAs and mutational burden can be very useful for prognosis and response to therapy (54).

To the best of our knowledge, this is the first time that our approach is used to partition a GB tumor tissue in its cellular components and provide its molecular profile. Single-cell CNA analysis has the potential to yield new insights into the molecular dynamics of cellular populations. Measuring single - cell genome alterations in tissues and cell populations will greatly advance clonal decomposition of malignant tissues, resolving rare cell populations genotypes and identifying DNA amplification and deletion states of individual cells, which are difficult to identify when cellular information is destroyed in bulk sequencing. A novel feature of our approach is also the capturing, by brightfield and immunofluorescence imaging, of morphologic features of cells permitting analytical integration with genomic properties. Moreover, the presence of double-stained cells in a framework of tumor cannibalism may represent specific tumor targets for future new strategies against cancer (46). Single-cell biology is leading to a new understanding of physiology and disease. The present resource of single-cell CNAs will permit new insights into genome heterogeneity, mutational processes, and clonal evolution in malignant tissues.

## FIGURE CAPTIONS

Figure 1S (Supplementary Data). Pie charts of the percentages of the double-stained cells and not stained cells found in GB01, GB02 and GB03.

## Bibliography

1. Stupp R, Hegi ME, Mason WP, Bent MJ van den, Taphoorn MJ, Janzer RC, Ludwin SK, Allgeier A, Fisher B, Belanger K, et al. Effects of radiotherapy with concomitant and adjuvant temozolomide versus radiotherapy alone on survival in glioblastoma in a randomised phase III study: 5-year analysis of the EORTC-NCIC trial. Lancet Oncol (2009) 10:459–466. doi:10.1016/S1470-2045(09)70025-7

2. Zhang X, Ding K, Wang J, Li X, Zhao P. Chemoresistance caused by the microenvironment of glioblastoma and the corresponding solutions. Biomed Pharmacother (2019) 109:39–46. doi:10.1016/J.BIOPHA.2018.10.063

3. Hambardzumyan D, Bergers G. Glioblastoma: Defining Tumor Niches. Trends in cancer (2015) 1:252. doi:10.1016/J.TRECAN.2015.10.009

4. Patel AP, Tirosh I, Trombetta JJ, Shalek AK, Gillespie SM, Wakimoto H, Cahill DP, Nahed B V., Curry WT, Martuza RL, et al. Single-cell RNA-seq highlights intratumoral heterogeneity in primary glioblastoma. Science (2014) 344:1396–1401. doi:10.1126/SCIENCE.1254257

5. Zhu Y, Parada LF. The molecular and genetic basis of neurological tumors. Nat Rev Cancer (2002) 2:616–626. doi:10.1038/NRC866

6. Yap TA, Gerlinger M, Futreal PA, Pusztai L, Swanton C. Intratumor Heterogeneity: Seeing the Wood for the Trees. Sci Transl Med (2012) 4: doi:10.1126/SCITRANSLMED.3003854

7. Meyer M, Reimand J, Lan X, Head R, Zhu X, Kushida M, Bayani J, Pressey JC, Lionel AC, Clarke ID, et al. Single-cell-derived clonal analysis of human glioblastoma links functional and genomic heterogeneity. Proc Natl Acad Sci U S A (2015) 112:851–856. doi:10.1073/PNAS.1320611111

8. Shlush LI, Hershkovitz D. Clonal evolution models of tumor heterogeneity. Am Soc Clin Oncol Educ book Am Soc Clin Oncol Annu Meet (2015) e662–e665. doi:10.14694/EDBOOK_AM.2015.35.E662

9. Tirosh I, Suvà ML. Dissecting human gliomas by single-cell RNA sequencing. Neuro Oncol (2018) 20:37. doi:10.1093/NEUONC/NOX126

10. Wang Y, Navin NE. Advances and Applications of Single-cell Sequencing Technologies. Mol Cell (2015) 58:598. doi:10.1016/J.MOLCEL.2015.05.005

11. Albertson DG, Collins C, McCormick F, Gray JW. Chromosome aberrations in solid tumors. Nat Genet (2003) 34:369–376. doi:10.1038/NG1215

12. Shlien A, Malkin D. Copy number variations and cancer. Genome Med (2009) 1: doi:10.1186/GM62

13. Stratton MR, Campbell PJ, Futreal PA. The cancer genome. Nature (2009) 458:719–724. doi:10.1038/NATURE07943

14. McLendon R, Friedman A, Bigner D, Van Meir EG, Brat DJ, Mastrogianakis GM, Olson JJ, Mikkelsen T, Lehman N, Aldape K, et al. Comprehensive genomic characterization defines human glioblastoma genes and core pathways. Nature (2008) 455:1061–1068. doi:10.1038/NATURE07385

15. Assem M, Sibenaller Z, Agarwal S, Al-Keilani MS, Alqudah MAY, Ryken TC. Enhancing Diagnosis, Prognosis, and Therapeutic Outcome Prediction of Gliomas Using Genomics. https://home.liebertpub.com/omi (2012) 16:113–122. doi:10.1089/OMI.2011.0031

16. Laks E, McPherson A, Zahn H, Lai D, Steif A, Brimhall J, Biele J, Wang B, Masud T, Ting J, et al. Clonal Decomposition and DNA Replication States Defined by Scaled Single-Cell Genome Sequencing. Cell (2019) 179:1207. doi:10.1016/J.CELL.2019.10.026

17. Adalsteinsson VA, Ha G, Freeman SS, Choudhury AD, Stover DG, Parsons HA, Gydush G, Reed SC, Rotem D, Rhoades J, et al. Scalable whole-exome sequencing of cell-free DNA reveals high concordance with metastatic tumors. Nat Commun 2017 81 (2017) 8:1–13. doi:10.1038/s41467-017-00965-y

18. Franch-Expósito S, Bassaganyas L, Vila-Casadesús M, Hernández-Illán E, Esteban-Fabró R, Díaz-Gay M, Lozano JJ, Castells A, Llovet JM, Castellví-Bel S, et al. CNApp, a tool for the quantification of copy number alterations and integrative analysis revealing clinical implications. Elife (2020) 9: doi:10.7554/ELIFE.50267

19. Fan F, Zhang H, Dai Z, Zhang Y, Xia Z, Cao H, Yang K, Hu S, Guo Y, Ding F, et al. A comprehensive prognostic signature for glioblastoma patients based on transcriptomics and single-cell sequencing. Cell Oncol (Dordr) (2021) 44:917–935. doi:10.1007/S13402-021-00612-1

20. A Classification of the Tumors of the Glioma Group on a Histogenetic Basis with a Correlated Study of Prognosis. J Am Med Assoc (1926) 87:268–268. doi:10.1001/JAMA.1926.02680040056039

21. Burrell RA, McGranahan N, Bartek J, Swanton C. The causes and consequences of genetic heterogeneity in cancer evolution. Nat 2013 5017467 (2013) 501:338–345. doi:10.1038/nature12625

22. Castro LNG, Tirosh I, Suvà ML. Decoding Cancer Biology One Cell at a Time. Cancer Discov (2021) 11:960. doi:10.1158/2159-8290.CD-20-1376

23. Tirosh I, Izar B, Prakadan SM, Wadsworth MH, Treacy D, Trombetta JJ, Rotem A, Rodman C, Lian C, Murphy G, et al. Dissecting the multicellular ecosystem of metastatic melanoma by single-cell RNA-seq. Science (2016) 352:189–196. doi:10.1126/SCIENCE.AAD0501

24. Taylor AM, Shih J, Ha G, Gao GF, Zhang X, Berger AC, Schumacher SE, Wang C, Hu H, Liu J, et al. Genomic and Functional Approaches to Understanding Cancer Aneuploidy. Cancer Cell (2018) 33:676. doi:10.1016/J.CCELL.2018.03.007

25. Reemst K, Noctor SC, Lucassen PJ, Hol EM. The Indispensable Roles of Microglia and Astrocytes during Brain Development. Front Hum Neurosci (2016) 10: doi:10.3389/FNHUM.2016.00566

26. Singh SK, Clarke ID, Terasaki M, Bonn VE, Hawkins C, Squire J, Dirks PB. Identification of a Cancer Stem Cell in Human Brain Tumors. Cancer Res (2003) 63:

27. Bao S, Wu Q, McLendon RE, Hao Y, Shi Q, Hjelmeland AB, Dewhirst MW, Bigner DD, Rich JN. Glioma stem cells promote radioresistance by preferential activation of the DNA damage response. Nat 2006 4447120 (2006) 444:756–760. doi:10.1038/nature05236

28. Lee JH, Lee JE, Kahng JY, Kim SH, Park JS, Yoon SJ, Um JY, Kim WK, Lee JK, Park J, et al. Human glioblastoma arises from subventricular zone cells with low-level driver mutations. Nat 2018 5607717 (2018) 560:243–247. doi:10.1038/s41586-018-0389-3

29. Matias D, Balça-Silva J, da Graça GC, Wanjiru CM, Macharia LW, Nascimento CP, Roque NR, Coelho-Aguiar JM, Pereira CM, Dos Santos MF, et al. Microglia/astrocytes– glioblastoma crosstalk: Crucial molecular mechanisms and microenvironmental factors. Front Cell Neurosci (2018) 12:235. doi:10.3389/FNCEL.2018.00235/BIBTEX

30. Gimple RC, Bhargava S, Dixit D, Rich JN. Glioblastoma stem cells: lessons from the tumor hierarchy in a lethal cancer. (2019) doi:10.1101/gad.324301

31. Keane L, Cheray M, Blomgren K, Joseph B. Multifaceted microglia - key players in primary brain tumour heterogeneity. Nat Rev Neurol (2021) 17:243–259. doi:10.1038/S41582-021-00463-2

32. Graeber MB, Scheithauer BW, Kreutzberg GW. Microglia in brain tumors. Glia (2002) 40:252–259. doi:10.1002/GLIA.10147

33. Leite DM, Zvar Baskovic B, Civita P, Neto C, Gumbleton M, Pilkington GJ. A human coculture cell model incorporating microglia supports glioblastoma growth and migration, and confers resistance to cytotoxics. FASEB J (2020) 34:1710–1727. doi:10.1096/FJ.201901858RR

34. Wolf SA, Boddeke HWGM, Kettenmann H. Microglia in Physiology and Disease. Annu Rev Physiol (2017) 79:619–643. doi:10.1146/ANNUREV-PHYSIOL-022516-034406

35. Bowman RL, Klemm F, Akkari L, Pyonteck SM, Sevenich L, Quail DF, Dhara S, Simpson K, Gardner EE, Iacobuzio-Donahue CA, et al. Macrophage Ontogeny Underlies Differences in Tumor-Specific Education in Brain Malignancies. Cell Rep (2016) 17:2445–2459. doi:10.1016/J.CELREP.2016.10.052

36. Maas SLN, Abels ER, Van De Haar LL, Zhang X, Morsett L, Sil S, Guedes J, Sen P, Prabhakar S, Hickman SE, et al. Glioblastoma hijacks microglial gene expression to support tumor growth. J Neuroinflammation 2020 171 (2020) 17:1–18. doi:10.1186/S12974-020-01797-2

37. Daneman R, Prat A. The blood–brain barrier. Cold Spring Harb Perspect Biol (2015) 7: doi:10.1101/CSHPERSPECT.A020412

38. Xie Y, He L, Lugano R, Zhang Y, Cao H, He Q, Chao M, Liu B, Cao Q, Wang J, et al. Key molecular alterations in endothelial cells in human glioblastoma uncovered through single-cell RNA sequencing. JCI Insight (2021) 6: doi:10.1172/JCI.INSIGHT.150861

39. Chouleur T, Tremblay ML, Bikfalvi A. Mechanisms of invasion in glioblastoma. Curr Opin Oncol (2020) 32:631–639. doi:10.1097/CCO.0000000000000679

40. Charalambous C, Hofman FM, Chen TC. Functional and phenotypic differences between glioblastoma multiforme-derived and normal human brain endothelial cells. J Neurosurg (2005) 102:699–705. doi:10.3171/JNS.2005.102.4.0699

41. Miebach S, Grau S, Hummel V, Rieckmann P, Tonn JC, Goldbrunner RH. Isolation and culture of microvascular endothelial cells from gliomas of different WHO grades. J Neurooncol (2006) 76:39–48. doi:10.1007/S11060-005-3674-6

42. Luissint AC, Artus C, Glacial F, Ganeshamoorthy K, Couraud PO. Tight junctions at the blood brain barrier: Physiological architecture and disease-associated dysregulation. Fluids Barriers CNS (2012) 9:1–12. doi:10.1186/2045-8118-9-23/TABLES/1

43. Wang R, Chadalavada K, Wilshire J, Kowalik U, Hovinga KE, Geber A, Fligelman B, Leversha M, Brennan C, Tabar V. Glioblastoma stem-like cells give rise to tumour endothelium. Nature (2010) 468:829–835. doi:10.1038/NATURE09624

44. Ricci-Vitiani L, Pallini R, Biffoni M, Todaro M, Invernici G, Cenci T, Maira G, Parati EA, Stassi G, Larocca LM, et al. Tumour vascularization via endothelial differentiation of glioblastoma stem-like cells. Nature (2010) 468:824–830. doi:10.1038/NATURE09557

45. Reduzzi C, Vismara M, Gerratana L, Silvestri M, De Braud F, Raspagliesi F, Verzoni E, Di Cosimo S, Locati LD, Cristofanilli M, et al. The curious phenomenon of dual-positive circulating cells: Longtime overlooked tumor cells. Semin Cancer Biol (2020) 60:344–350. doi:10.1016/J.SEMCANCER.2019.10.008

46. Fais S. Cannibalism: a way to feed on metastatic tumors. Cancer Lett (2007) 258:155–164. doi:10.1016/J.CANLET.2007.09.014

47. Coopman PJ, Do MT, Thompson EW, Mueller SC. Phagocytosis of cross-linked gelatin matrix by human breast carcinoma cells correlates with their invasive capacity. Clin Cancer Res (1998) 4:

48. Persson A, Englund E. The glioma cell edge--winning by engulfing the enemyã Med Hypotheses (2009) 73:336–337. doi:10.1016/J.MEHY.2009.03.042

49. Chang GHF, Barbaro NM, Pieper RO. Phosphatidylserine-dependent phagocytosis of apoptotic glioma cells by normal human microglia, astrocytes, and glioma cells. Neuro Oncol (2000) 2:174–183. doi:10.1093/NEUONC/2.3.174

50. Huysentruyt LC, Seyfried TN. Perspectives on the mesenchymal origin of metastatic cancer. Cancer Metastasis Rev (2010) 29:695–707. doi:10.1007/S10555-010-9254-Z

51. Wilhelmsson U, Eliasson C, Bjerkvig R, Pekny M. Loss of GFAP expression in high-grade astrocytomas does not contribute to tumor development or progression. Oncogene (2003) 22:3407–3411. doi:10.1038/sj.onc.1206372

52. Hieronymus H, Murali R, Tin A, Yadav K, Abida W, Moller H, Berney D, Scher H, Carver B, Scardino P, et al. Tumor copy number alteration burden is a pan-cancer prognostic factor associated with recurrence and death. Elife (2018) 7: doi:10.7554/ELIFE.37294

53. Gómez-Rueda H, Martínez-Ledesma E, Martínez-Torteya A, Palacios-Corona R, Trevino V. Integration and comparison of different genomic data for outcome prediction in cancer. BioData Min (2015) 8: doi:10.1186/S13040-015-0065-1

54. Roh W, Chen PL, Reuben A, Spencer CN, Prieto PA, Miller JP, Gopalakrishnan V, Wang F, Cooper ZA, Reddy SM, et al. Integrated molecular analysis of tumor biopsies on sequential CTLA-4 and PD-1 blockade reveals markers of response and resistance. Sci Transl Med (2017) 9: doi:10.1126/SCITRANSLMED.AAH3560

