## Supplementary figures and images for "Single-cell molecular characterization to partition the human glioblastoma tumor microenvironment (TME) genetic background"

### Figure 1S

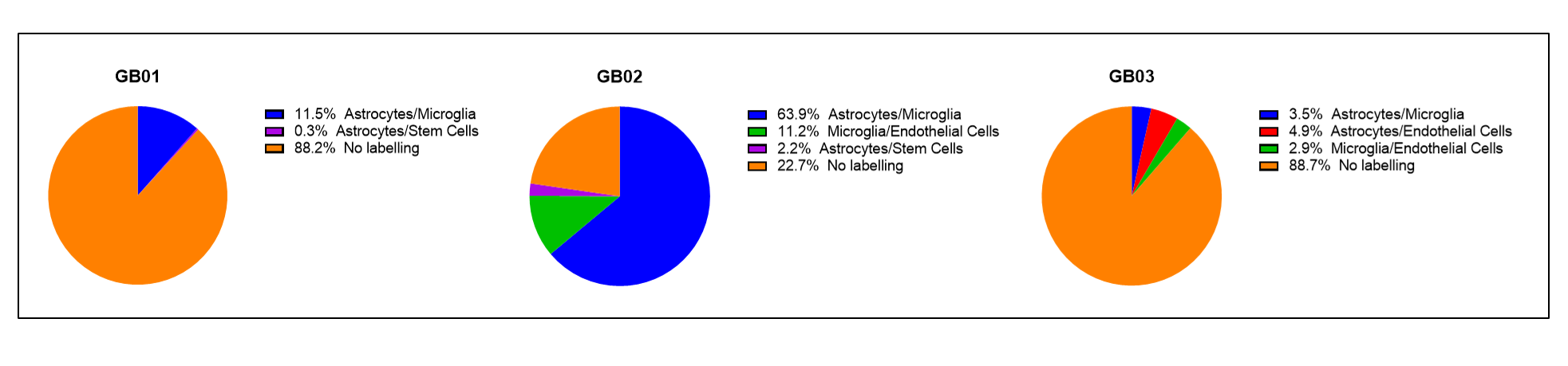
